# Relationship between extreme species richness and Holocene persistence of forest-steppe grasslands in Transylvania, Romania

**DOI:** 10.1101/2024.01.23.576840

**Authors:** Jan Novák, Pavel Šamonil, Jan Roleček

## Abstract

The most species-rich grasslands worldwide are known from the Carpathian Mts and their periphery in East-Central Europe. They occur in forest-steppe regions, transitional between temperate forest and arid steppe biomes. Their climate, largely suitable for forests, raises questions about the origin of these grasslands. Have they been forested in the past, or locally maintained through a disturbance regime? We addressed these questions to contribute to the broader understanding of Holocene dynamics of open habitats in temperate Europe. We employed soil charcoal analysis and soil morphology to reconstruct past representation of woody species with fine spatial resolution. Our study area was Romanian Transylvania, a region renowned for a well-developed forest-steppe. Six soil profiles along a climatic gradient were assessed: four in forest-steppe grasslands, two in grasslands in adjacent forest region. The results revealed profound differences between forest-steppe and forest grasslands. Forest-steppe profiles showed Phaeozems with low specific anthracomass and continuous dominance by *Juniperus*, suggesting a long-term presence of grasslands. Forest profiles showed Luvisols with higher anthracomass and abundant charcoal of broad-leaved trees, indicating establishment after deforestation. The high radiocarbon ages of charcoals in basal soil horizons point to a glacial origin of soils and the link of forest-steppe grasslands to glacial forests. Siberian hemiboreal forests and related grasslands may be modern analogues of the reconstructed ecosystems, sharing many species with present day forest-steppe. We highlight the role of disturbances such as fire, herbivore grazing, and human activities in shaping the forest-steppe over time, contributing to the formation of today’s richest grasslands.

## Introduction

Forest-steppes are diverse landscape mosaics of variously dense forests and steppe grasslands that occur in continental regions, at the transition between temperate forest and arid steppe biomes. They are widespread in Eurasia (Dokhman, 1968; Erdős et al., 2018), but similar ecosystems occur also in North America and other continents, where the term forest-steppe is not widely used for them (Billings and Barbour, 2000; Myster, 2012). From the point of view of vegetation composition, forest-steppes differ from other landscape mosaics comprising grasslands and forests (e.g. mountain wood pastures and urban parks) by the abundance of steppe grasses and herbs.

According to current knowledge, the most species-rich forest-steppe grasslands occur in the Carpathian region and its periphery in East-Central Europe (Biurrun et al., 2021; Roleček et al., 2021; Wilson et al., 2012). It is striking that these communities are confined to relatively productive habitats with conditions suitable for shady mesophilous forests, dominated mostly by *Quercus*-*Carpinus* or *Fagus* (Dengler et al., 2012; Roleček et al., 2014; Soó, 1927). The occurrence of open and semi-open ecosystems in potentially forested landscapes raises questions about their origin and long-term dynamics. In addition to climate, other possible driving factors such as topography and especially the disturbance regime come to the fore (Chytrý et al., 2022; Erdős et al., 2022; Sümegi et al., 2012).

Addressing these questions using a traditional palaeoecological approach is not straightforward. The pollen record, as the most common palaeoecological proxy (Birks, 2019), may mix signals from different components of the forest-steppe mosaics due to its relatively low spatial resolution, especially in the case of records from large open sites such as lakes and peatlands (Prentice, 1985; Bradshaw, 1991). In addition, the quantitative representation of species in the pollen record is known to be biased (Andersen, 1967; Bunting et al., 2013; Kuneš et al., 2019), which may also hinder the reconstruction of habitat mosaics. Although the application of numerical tools may alleviate such biases (Prentice, 1985; Sugita, 2007), the question of the past distribution of forests and open habitats in the forest-steppe remains. Here, a proxy recording local development with fine spatial resolution may be a useful tool.

In the present study we focused on the long-term development of the most species-rich forest-steppe grasslands in Romanian Transylvania. This region is renowned for a well-developed forest-steppe, at least partly determined by its relatively dry climate and rugged topography (Bădărău, 2005; Kun et al., 2004; Varga et al., 2000). Palaeoecological studies have brought evidence of Holocene continuity of open habitats in the region (Feurdean et al., 2015, 2018). However, the most species-rich grasslands occur on the periphery of the Transylvanian Basin (Dengler et al., 2012; Roleček et al. 2021), in an area that is climatically favourable for mesophilous forests; such forests do indeed occur near the well-preserved grassland sites (Bădărău, 2005). Also local soils show signs of historical occurrence of forests or at least woody species (Püntener et al., 2023). They are no longer Chernozems as in the central part of the Transylvanian Basin, but Phaeozems and even Luvisols, i.e. soils dominated by the vertical translocation of substances in the environment of the transition between Ustic and Udic moisture regime (Eckmeier et al., 2007; Püntener et al., 2023).

Thus, two alternative scenarios for the long-term development of the extremely species-rich grasslands in the Transylvanian Basin can be put forward, which both find support in published studies: i) Either the present grasslands were forested in the past, most likely during the Middle Holocene, when the climate was particularly suitable for the spread of forests and human impact was still relatively low; in this case non-forest species appeared here mainly through later immigration from other parts of the Transylvania where the climate, habitat conditions or disturbance regime were more suitable for their persistence; (ii) Or the open character of the sites where species-rich steppe grasslands occur today was locally maintained throughout the Holocene; this probably could not have been achieved other than by a specific long-term disturbance regime.

To test these hypotheses, we employed soil charcoal analysis as a tool to reconstruct past representation of woody species under fire disturbance with fine spatial resolution. At the same time, we studied the soil morphology, which also reflects the historical development of the landscape. Elucidating these issues may also contribute to answering the more general question of how much forest, as a competitively superior community, was present on mesic sites during the postglacial development of vegetation in the lowlands of temperate Europe.

### Study area

All the study sites are situated in northwestern part of the Transylvanian Basin near the city of Cluj (Figure 1, Table 1): four of them in the forest-steppe region of the Transylvanian Plain (Valea Florilor, Boj-Cătun) and Someş Plateau (Fânaţele Clujului I, II), the other two in the adjacent forest regions of the Feleacu Massif (Sălicea) and Hășdate-Vlaha Depression (Lita) (Rosean, 2020). The sites are found on an altitudinal and climatic gradient, with altitude and precipitation decreasing and temperatures increasing approximately from west to east, with some local variations related to topography (Table 1). The irregular spacing of sites follows the distribution of species-rich steppe grasslands, which were the focus of our sampling.

**Figure 1.**
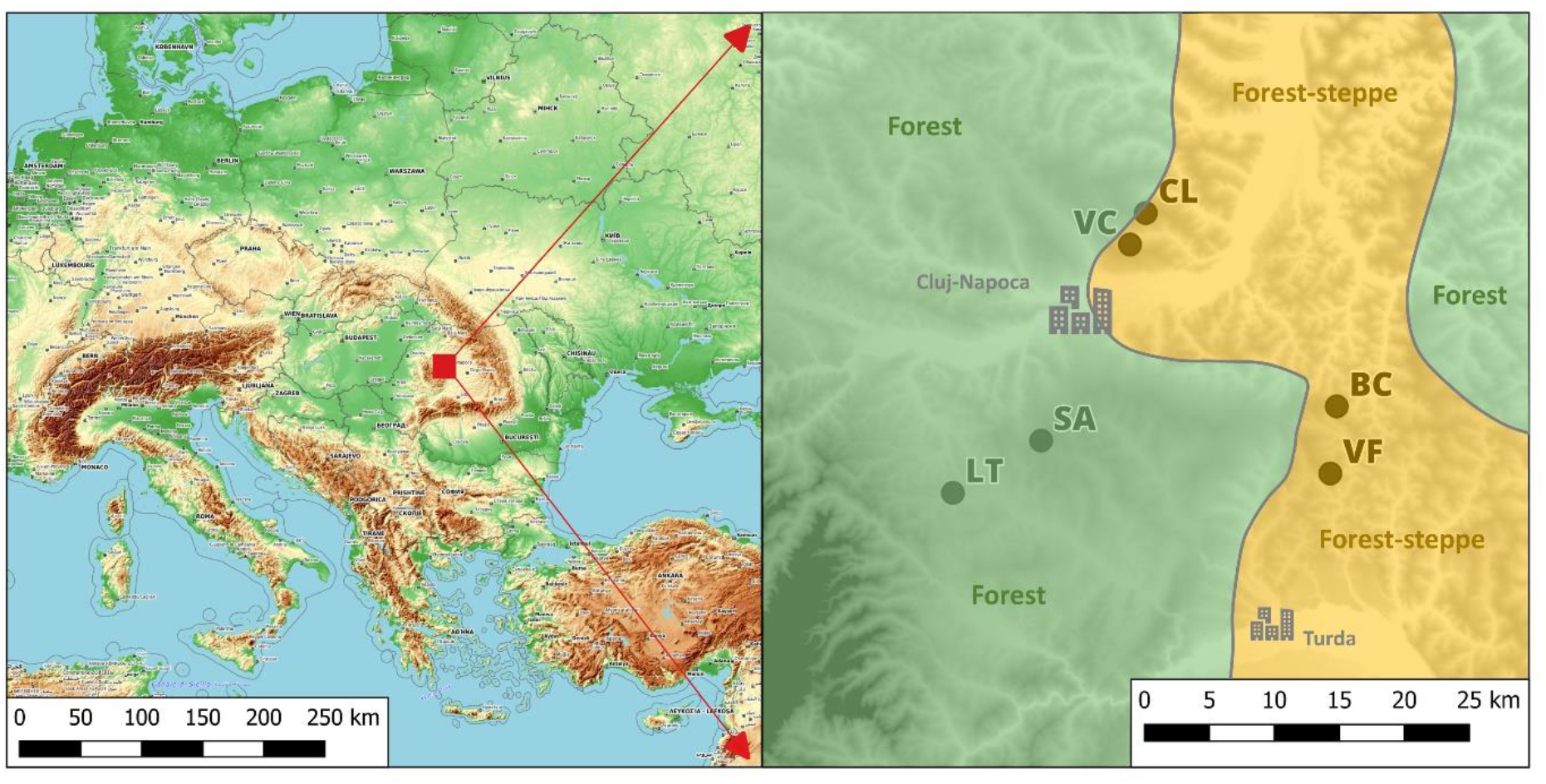
Map of the study area. Delimitation of the forest-steppe region adapted from Niklfeld (1974) and Varga et al. (2000). Black dots indicate study sites (see Table 1 for an explanation of the study site codes).

**Table 1.** Basic physiographic characteristics of the study sites. Climatic data taken from CHELSA V2.1 (Karger et al., 2017). Delimitation of the forest-steppe region follows Niklfeld (1974) and Varga et al. (2000).

|  | Site |  | Latitude | Longitude | Altitude | Annual Mean | Annual |
| --- | --- | --- | --- | --- | --- | --- | --- |
| Site | code | Region | [°N] | [°E] | [m a.s.l.] | Temperature | Precipitation |
| Valea Florilor | VF | Forest-steppe | 46.660765 | 23.841105 | 450 | 9.1 | 589 |
| Boj-Cătun | BC | Forest-steppe | 46.706936 | 23.847622 | 410 | 9.1 | 585 |
| Fânațele Clujului I<br>-Valea lui Craiu | VLC | Forest-steppe | 46.8396 | 23.6563 | 550 | 8.4 | 683 |
| Fânațele Clujului II<br>-Valea Caldă | VC | Forest-steppe | 46.818059 | 23.640519 | 430 | 8.5 | 645 |
| Lita | LT | Forest | 46.64745 | 23.463101 | 605 | 8.1 | 703 |
| Sălicea | SA | Forest | 46.683417 | 23.551528 | 785 | 7.0 | 747 |

The dominant soil-forming process is the gradual translocation of clay and organic matter (Püntener et al., 2023; own data). The bedrock is in all cases Tertiary sediments, mainly calcareous claystones and marlstones, with a higher proportion of sand at Sălicea. From a topographical point of view, the research focused on the upper to middle parts of the slopes, we avoided visible surface disturbance due to landslides, debris flow or slumps during site selection. However, we accepted and assumed colluviation and the presence of creep. These processes were confirmed radiometrically in the profiles (Püntener et al., 2023). Moreover, the radiometry revealed a limited occurrence of past mechanical disturbance at some sites, e.g. in the form of ploughing, which could not be detected in the field.

Mesic forest-steppe meadows of the *Brachypodio-Molinietum* association are the dominant grassland type at Boj-Cătun and Fânaţele Clujului II sites, while slightly drier association *Stipetum tirsae* prevails at Valea Florilor and Fânaţele Clujului I (Willner et al., 2019). Less species-rich semi-dry grassland of the *Cirsio-Brachypodion* alliance dominates at Lita site, while more acidophilous, previously grazed grasslands with some steppe elements (*Cynosurion cristati* alliance) prevails at Sălicea site. The potential forest vegetation includes mainly *Quercus*-dominated forests in the case of the driest part of the study area (Boj-Cătun, Valea Florilor), *Quercus-* and *Carpinus*-dominated forests in the case of the mid-altitude sites (Fânaţele Clujului I, II, Lita) and *Carpinus-* and *Fagus*-dominated forests in the case of the highest site Sălicea. The current landscape around all four forest-steppe sites is almost devoid of forests, the main exception being small plantations of *Robinia pseudoacacia*.

## Materials and methods

### Pedoanthracological analysis

We employed soil charcoal analysis to provide a record of past local development with fine spatial resolution (e.g., Clark et al., 1988; Figueiral and Terral, 2002; Heinz et al., 2004). In general, charcoal fragments originate from at least one past fire event and are essentially ubiquitous in soils (Robin et al., 2013). The quality of the soil charcoal record depends on the topography of the source area, which is usually quite small, so charcoal can be used to reconstruct vegetation dynamics at a local scale (Robin et al., 2013; Touflan et al., 2010).

We analysed charcoal fragments from six soil profiles which were taken in extensively managed grasslands along a climatic gradient. Four profiles were deliberately placed at some of the best-preserved stands of extremely species-rich forest-steppe grasslands in a forest-steppe region, other two in grasslands with lower species richness in adjacent forest region, where we assumed a longer-term presence of forest in the past. We refer to the latter as forest grasslands. The standard soil profiles were excavated by hand and 10 litres of soil were taken from each regularly spaced 10 cm thick layer. The soils were pedomorphologically evaluated (Schoeneberger et al., 2012) and the soils were classified according to the international soil taxonomy (IUSS Working Group WRB, 2015). The extracted samples were dried (60 °C until completely dry) and weighed in the lab. Charcoal fragments were extracted using water flotation and a wet-sieving procedure (Carcaillet and Thinon, 1996) with a minimum mesh size of 0.4 mm. The sieved fraction was then dried again; charcoal fragments were hand-picked under a binocular lens and weighed.

Taxonomic determination was performed on all charcoal fragments larger than 1 mm, except for a single sample with the highest charcoal quantity, which was subsampled. Determination is usually possible at the genus or sometimes species level, successful determination of fragments smaller than 1 mm is much less likely or impossible (Robin et al., 2013). *Cornus*, *Prunus* and *Rosa* are referred to as “shrubs” in the figures. The small charcoals of *Picea* and *Larix* are difficult to distinguish and the two taxa are therefore lumped together and referred to as *Picea*/*Larix*.

Determination was performed on a reflected light microscope (Jenatech Inspection, Carl Zeiss Jena; magnification 200–500×) and a binocular lens (Leica S9E; magnification 40–220×) using a reference collection and available identification keys (Benkova and Schweingruber, 2004; Schoch et al., 2004; Schweingruber, 1978). Determined charcoal fragments were weighed to an accuracy of the nearest 0.1 mg. Charcoal concentration was expressed as specific anthracomass (mg of charcoal per kg of soil; SA) and was calculated per layer (SAL), profile (SAP) or specific taxon within a profile (Talon, 2010).

A total of 26 charcoal samples (with a minimum of three and an average of four per profile) were selected for radiocarbon dating to obtain an approximate time frame for each profile. AMS dating was performed in two laboratories: Center for Applied Isotope Studies, University of Georgia, US (23 dates) and Isotoptech Zrt., Debrecen, Hungary (three dates). For details, see Table 2. All dates were calibrated using the *rintcal* package in R software version 4.3.1 (R Core Team, 2023) and median values of the resulting probability intervals were used further in the text.

**Table 2.** List of 26 charcoal radiocarbon dates. Median values rounded to units are reported for calibrated age (see the text for details).

| Site code | Depth | Age (BP) | Range | Calibrated |  | Dated taxon |
| --- | --- | --- | --- | --- | --- | --- |
|  |  |  |  | age | Lab code |  |
| VF | 30 | 630 | ±20 | 594 | UGAMS-59063 | *Juniperus* |
| VF | 50 | 4050 | ±25 | 4516 | UGAMS-59064 | *Juniperus* |
| VF | 70 | 1110 | ±20 | 1005 | UGAMS-62554 | *Stipa* |
| VF | 100 | 7520 | ±30 | 8347 | UGAMS-59065 | *Juniperus* |
| VF | 160 | 40590 | ±240 | 43645 | UGAMS-59066 | *Larix* |
| VF | 180 | 11666 | ±71 | 13522 | DeA-40575 | *Juniperus* |
| BC | 40 | 5250 | ±25 | 5994 | UGAMS-62555 | *Juniperus* |
| BC | 60 | 12150 | ±40 | 14052 | UGAMS-62556 | *Pomoideae* |
| BC | 90 | 12160 | ±40 | 14061 | UGAMS-62557 | *Juniperus* |
| BC | 120 | 12540 | ±60 | 14850 | UGAMS-62558 | *Juniperus* |
| BC | 160 | 19140 | ±50 | 23025 | UGAMS-62559 | *Juniperus* |
| VLC | 60 | 4140 | ±25 | 4684 | UGAMS-56033 | *Juniperus* |
| VLC | 80 | 11870 | ±50 | 13700 | UGAMS-59059 | *Juniperus* |
| VLC | 100 | 9620 | ±30 | 10937 | UGAMS-56034 | *Juniperus* |
| VC | 40 | 4610 | ±30 | 5401 | UGAMS-59054 | *Juniperus* |
| VC | 60 | 1820 | ±25 | 1718 | UGAMS-59055 | *Fraxinus* |
| VC | 90 | 3280 | ±20 | 3485 | UGAMS-59056 | *Pomoideae* |
| VC | 120 | 14510 | ±50 | 17679 | UGAMS-59057 | *Juniperus* |
| VC | 140 | 11360 | ±30 | 13235 | UGAMS-59058 | *Juniperus* |
| LT | 30 | 2131 | ±28 | 2098.6 | DeA-40572 | *Quercus* |
| LT | 60 | 3940 | ±25 | 4382 | UGAMS-59060 | *Quercus* |
| LT | 90 | 2870 | ±25 | 2990.4 | UGAMS-59061 | *Quercus* |
| LT | 120 | 590 | ±25 | 603.3 | UGAMS-59062 | *Quercus* |
| SA | 30 | 62 | $\pm 24$ | Modern | DeA-40580 | *Quercus* |
| SA | 60 | 70 | ±n.d. | Modern | UGAMS-62551 | *Quercus* |
| SA | 180 | 11500 | $\pm 50$ | 13374.9 | UGAMS-62552 | *Juniperus* |

### Taxonomic concept

Taxonomic concept and nomenclature of vascular plants follow Euro+Med plantbase (Euro+Med 2006+), with the exception of broadly conceived taxa *Brachypodium pinnatum* s.lat. (= *B. pinnatum* + *B. rupestre*), *Poa pratensis* s.lat. (= *P. angustifolia* + *P. pratensis*), *Pulmonaria mollis* s.lat. (= *P. dacica* + *P. mollis*), and *Vicia cracca* s.lat. (= *V. cracca* + *V. tenuifolia*).

### Data analysis

The taxon-by-sample matrix with log-transformed abundances was subjected to the detrended correspondence analysis (DCA) in Canoco 5 software (Šmilauer and Lepš, 2014) to find the main gradients in the taxonomic composition of charcoal samples. The abundance values of rare species were downweighted according to the Canoco 5 protocol. The first gradient was 4.9 SD units long, so the unimodal method was appropriate. To place the ordination results in a time frame, we fitted the available charcoal radiocarbon age values with a generalized additive model and projected them onto the species ordination plot as isolines. We further grouped charcoal samples by sites, connected the samples of each sample series with lines, and projected the resulting series onto an ordination plot to illustrate taxonomic turnover along the profiles. The statistical relationship between charcoal age and depth at which it was found was evaluated with multiple regression analysis using the *stats* package in R software version 4.3.1 (R Core Team, 2023).

## Results

### Charcoal taxonomic composition and anthracomass

In total, we analysed 90 samples from six soil profiles, which resulted in 2742 individual identifications. We recorded 17 woody taxa in the whole charcoal assemblage: *Abies, Acer, Betula, Carpinus, Cornus*, *Fagus, Fraxinus, Juniperus, Larix/Picea, Pinus,* Pomoideae*, Prunus, Quercus, Populus, Rosa, Tilia* and *Ulmus*. Values of specific anthracomass per layer (SAL) were low, with a median value of 2.2 mg/kg, and varied widely between 0.2 and 898.5 mg/kg. Maximum values were recorded in the uppermost layer of the Sălicea profile, which was probably of recent age based on the results of radiocarbon dating.

We recorded notable differences in charcoal taxonomic composition and SAL values between species-rich forest-steppe grasslands and forest grasslands (Figure 2). Forest-steppe grasslands were distinguished by low SAL (0.2–36.5 mg/kg, median 1.55 mg/kg), while forest grasslands showed significantly higher values (0.4–895.5 mg/kg, median 17.45 mg/kg).

**Figure 2.**
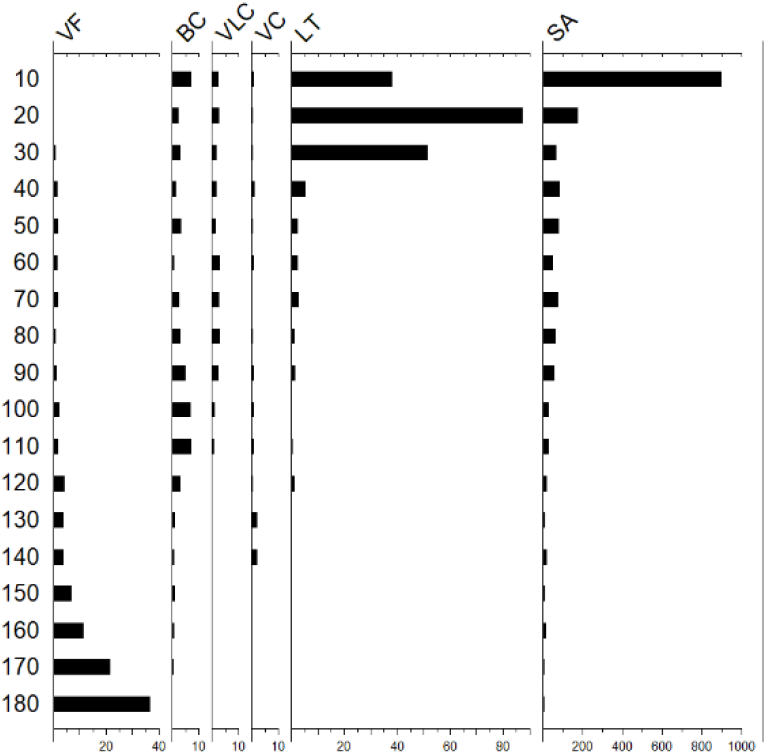
Values of specific anthracomass per layer (SAL, mg charcoal/kg sediment) in the analysed soil profiles.

### 4. 2. Differences between sites and temporal dynamics

A highly significant positive relationship between charcoal age and depth was found across profiles (R^2^ adj. = 0.49, p = 0.0001).

Pedoanthracological results clearly document different composition of charcoal assemblages of species-rich forest-steppe grasslands and forest grasslands (Figure 3, 4). Species composition of both vegetation types during Glacial and Late Glacial period was more or less similar. The dominance of *Juniperus* charcoals and common presence of *Pinus* was recorded in all analysed profiles with the exception of the profile Lita, which apparently recorded only Holocene history according to radiocarbon dating. Profiles Valea Florilor and Sălicea stood out with the presence of *Larix/Picea* charcoals. The highest SAL values in the Valea Florilor profile (Figure 2) were recorded during the Glacial period, suggesting most abundant tree presence at the site during this period.

**Figure 3.**
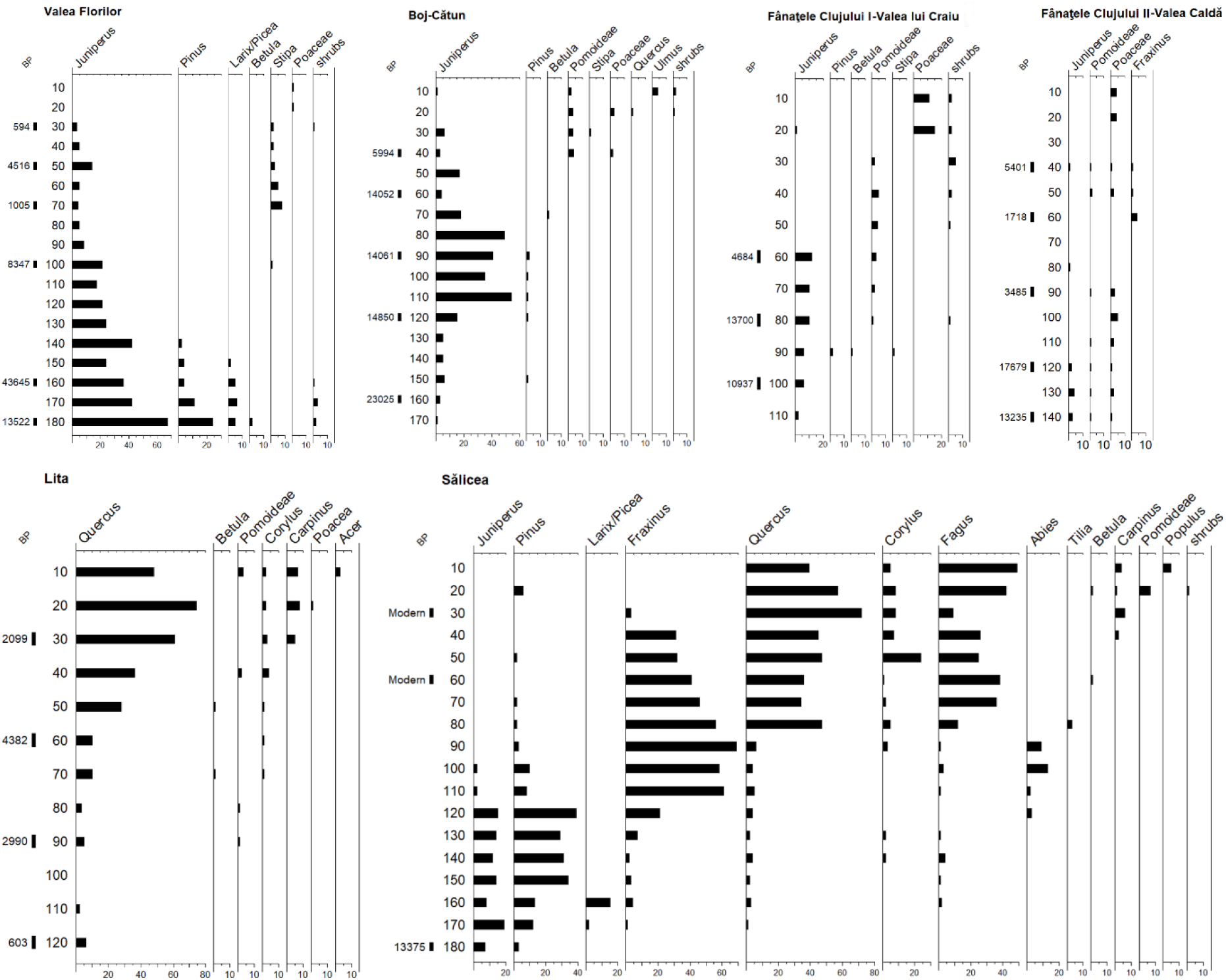
Taxonomic composition of the analysed charcoal assemblages. Numbers of charcoals exceeding 1 mm in size and radiocarbon dates of selected charcoals are given (see Table 2 for details on C14 ages).

Significant differences in charcoals composition between forest-steppe and forest grasslands have been recorded for the Holocene period. The species composition of the forest-steppe profiles was continuously dominated by *Juniperus* until the Late Holocene, with other heliophilous woody taxa (Pomoideae*, Prunus, Rosa*) admixed. Deciduous broad-leaved trees (*Ulmus, Quercus* or *Fraxinus*) were recorded only occasionally and with low abundance. Charred owns of the steppe grass *Stipa* and non-specific macroremains of Poaceae were also revealed. SAL values of all forest-steppe profiles were very low during the Holocene (Figure 2).

Holocene parts of pedoanthracological profiles of forest grasslands were distinguished by the abundance of deciduous broad-leaved trees (*Quercus, Carpinus* and *Corylus* in both profiles, plus *Fraxinus* and *Fagus* at Sălicea) (Figure 3). A number of other woody taxa (*Abies*, *Acer, Betula, Cornus,* Pomoideae and *Populus*) were recorded with lower abundance. The topsoil layers were characterized by higher SAL values (Figure 2).

The soils were classified as Chernic (Luvic) Phaeozems in drier lower elevations of forest-steppes (Valea Florilor, Boj-Cătun, Fânaţele Clujului I, II) and as Stagnic Luvisols in wetter higher elevations (Lita, Sălicea). While the forest-grassland soils were in an advanced stage of clay translocation, associated with the formation of the eluvial E horizon, the forest-steppe soils had not yet developed a metamorphic B horizon. Even in Phaeozems, the initial translocation of organic matter and clay has been observed morphologically as clay and organic matter coatings on soil aggregates.

The DCA plots (Figure 4) illustrate the divergent dynamics of forest-steppe and forest grasslands since the Late Glacial. Particularly instructive are the deep profiles of Valea Florilor (forest-steppe region) and Sălicea (forest region).

**Figure 4.**
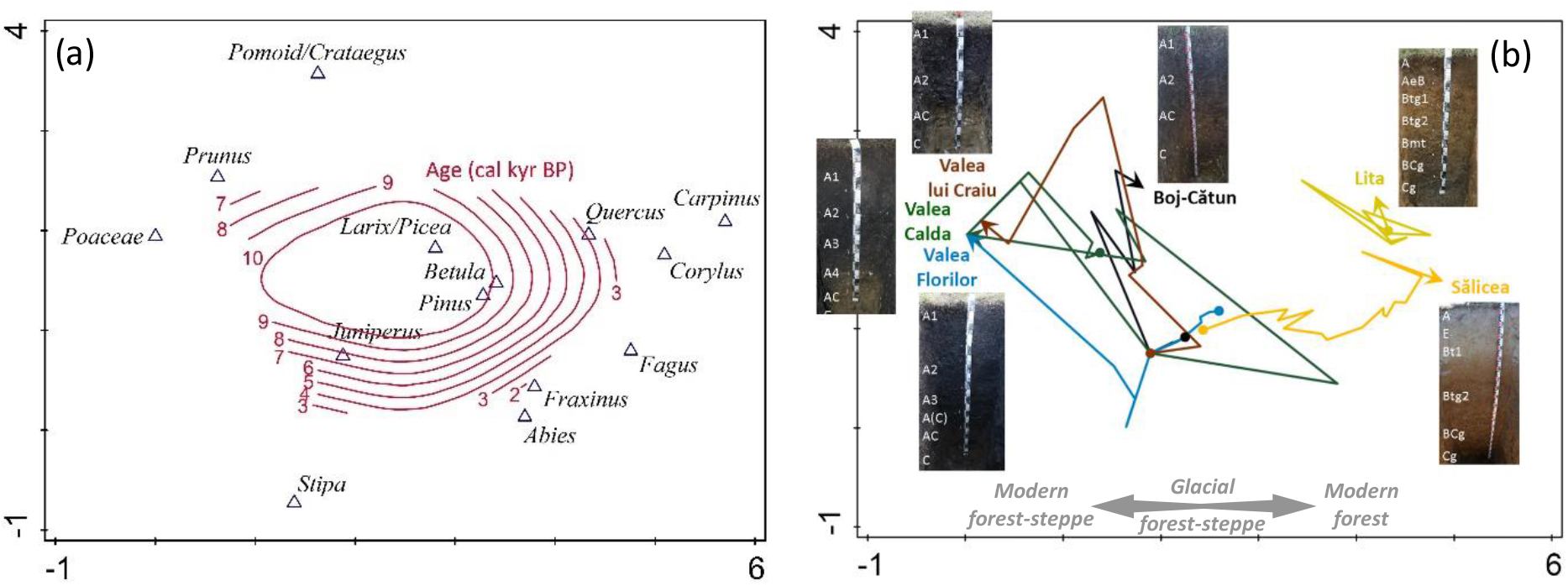
Results of DCA of taxonomic composition of individual charcoal samples. Plot (a, left) shows the centroids of each taxon (in italics) along the two main compositional gradients, with the available charcoal radiocarbon age values fitted by a generalized additive model and projected onto the ordination plot as isolines. Species with less than three occurrences in the dataset were omitted. Plot (b, right) shows sample series illustrating the taxonomic turnover in each analysed soil profile from the bottom layer (indicated by a dot) to the topsoil (indicated by an arrow and the site name).

## Discussion

The analyzed pedoanthracological profiles were placed along an altitudinal and climatic gradient running from the core of the forest-steppe area between Cluj and Turda, rich in dry steppe grasslands and saline habitats, to forested uplands rich in mesic grasslands and broad-leaved forests dominated by *Quercus-Carpinus* and *Fagus*. A common feature of all study sites was the presence of semi-dry grassland vegetation, and a focus on species-rich forest-steppe meadows (*Brachypodio-Molinietum* and *Stipetum tirsae* associations; Willner et al., 2019) determined the selection of sampling sites. At present, this vegetation is rare in the region under study, mainly due to intensive agriculture. The species composition of grasslands is also affected by abandonment, especially in the forest region, and by intensification of grazing in other places (Bădărău, 2005; Roleček et al., 2021). For this reason, little disturbed sites with an adequate species composition, suitable for pedoanthracological research, are scarce in the territory.

Despite the limited number of profiles assessed, soil charcoal analysis yielded multiple results relevant for reconstructing the history of the Transylvanian forest-steppe grasslands. Surprisingly, we gained insights not only into their Holocene development, but also into their deeper past, so that we can further our understanding of their ancestral ecosystems in the Last Glacial period. The exceptionally high ^14^C ages of charcoal fragments in the basal soil horizons indicate a surprisingly early origin of local soils. The age of chernozem-like soils was previously commonly placed as late as the Holocene (Eckmeier et al., 2007); however, some of the soils in Transylvania are obviously of glacial age (Pendea et al., 2002), which is also supported by our results of the radiometric analysis of erosion processes (Püntener et al., 2023).

### Contrasting Holocene development of grasslands in the forest-steppe and forest zones

Our results clearly support the hypothesis of the Holocene continuity of species-rich forest-steppe grasslands on mesic sites in Transylvania. A striking feature of all four forest-steppe profiles analysed was the absence of charcoals of temperate trees. The only exceptions are a small amount of *Fraxinus* charcoal dated to the Roman period (1720 cal kyr BP) at the Fânaţele Clujului II-Valea Caldă site and the charcoal fragments of *Quercus* and *Ulmus* in the top 20 cm of the profile at the Boj-Cătun site, likely of recent origin. This pattern sharply contrasts with the abundance and diversity of charcoal of temperate (nemoral) trees and shrubs (*Abies*, *Acer*, *Carpinus*, *Cornus*, *Corylus*, *Fagus*, *Fraxinus*, *Quercus*, *Tilia*) in both reference profiles, situated in the forest grasslands. The differences between forest-steppe and forest grasslands profiles are also clearly indicated by significantly different SAL values and topsoil (A horizon) depths.

Translocation processes in soils were significantly more visible in the control forest grasslands than in the forest-steppes. Due to the more humid, colder climate (Lapenis, et al. 2008), and the strengthening influence of the forest vegetation (Hubová, et al. 2018; Smolíková, 1967), the forest-grassland soils, Luvisols, showed a clearly developed metamorphic illuvial argic Bt horizon and at the same time an eluvial E horizon. On the other hand, forest-steppe soils, Phaeozems, were in an initial stage of (e.g. clay) translocation: clay coatings were practically absent in the soils of the Fânaţele Clujului I and II sites (Püntener et al. 2023). This supports the idea of a long-term local presence of grasslands, as grasses may slow down the progressive development of soils (pH decrease, leaching of carbonates, translocation of nutrients and clay; Eckmeier, et al. 2007; Marković, et al. 2018). Also, the exceptionally thick upper mineral mollic A horizon (IUSS Working Group WRB, 2022) in these soils, which is rich in organic carbon (local organic C concentrations reach up to 10%), is traditionally considered to result from steppe history (Eckmeier et al. 2007). The integration of knowledge on vegetation development, disturbance regime and soil evolution within the forest-steppes, including feedback interactions, requires further study.

We interpret the described differences as a consequence of the different Holocene history of the grasslands in the two regions: while the forest-steppe grasslands are ancient and have never been completely overgrown by forest, the grasslands in the forest zone are younger and were established after deforestation. The successional instability of the forest-steppe grasslands is only manifested by the regular occurrence of shrubs (mainly Pomoideae and *Juniperus*; for the latter species see discussion below). The sparse occurrence of woody plants is also indicated by the long-term low SAL values. Such low values have rarely been reported from Europe, one exception being the forest-steppe foothills of the White Carpathians (Novák et al., 2019). Although SAL values are low, we consider fires to be an important environmental factor affecting the character of grassland vegetation, especially in the last six thousand years (Figure 5). The two different histories of the regions are also reflected in different soil-taxonomic units reflecting landscape memory, i.e. Phaeozems on the one side and Luvisols on the other.

**Figure 5.**
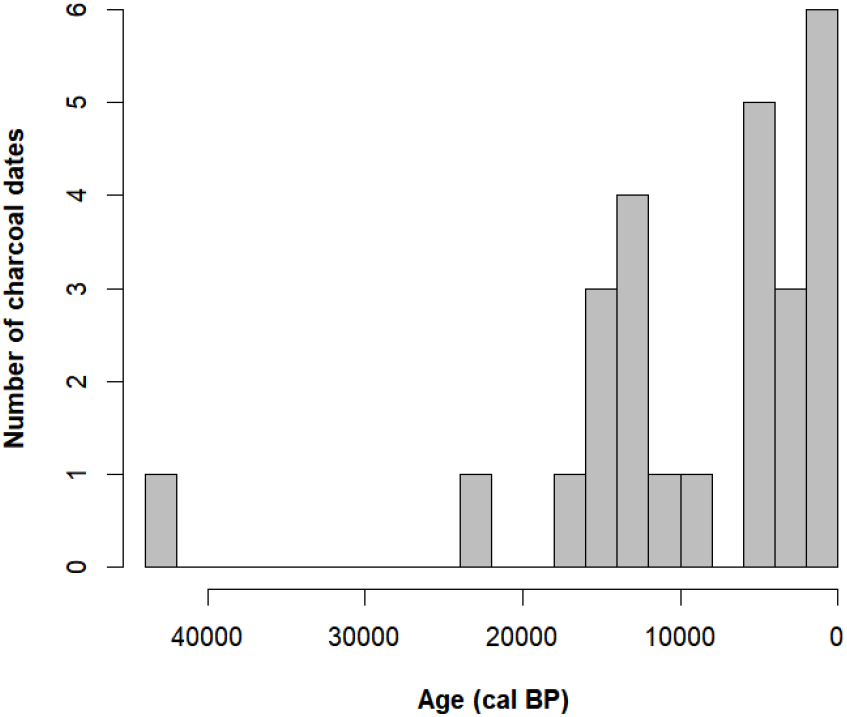
Frequency distribution of 26 calibrated radiocarbon dates obtained from soil charcoals in 2,000-year intervals.

### Further evidence

Our results are consistent with the multi-proxy study of the Transylvanian forest-steppe by Feurdean et al. (2015). They estimated the proportion of open habitats in the Middle Holocene, i.e. the period suitable for forest expansion and critical for grassland persistence, to be about 35% according to REVEALS model. Our results then suggest that open habitats were not confined to extreme sites such as steep southern slopes, saline soils and lakes, but persisted also on some mesic sites. This agrees also with the biogeographical evidence: Transylvanian grasslands harbour a group of rare mesophilous species with isolated occurrences (e.g. *Adenophora liliifolia*, *Adonis volgensis*, *Colchicum bulbocodium*, *Paeonia tenuifolia* and *Serratula coronata*) that could hardly persist on ecologically extreme sites (Bădărău, 2005; Roleček et al., 2014). As poor dispersers, they are likely relicts of once more widespread open ecosystems. Mesic grasslands also seem to be favoured by some animal species of open habitats that have isolated populations in the Transylvanian forest-steppe, such as *Poecilimon intermedius* (Togănel, 2000), *Vipera ursinii* (Krecsák and Zamfirescu, 2008) and *Sicista subtilis* (Cserkész et al., 2016). ecologically. They may also indicate ancient origin of local grasslands.

It is worth noting that a number of isolated occurrences of rare light-demanding plant species of mesic habitats (e.g. *Achillea impatiens*, *Adenophora liliifolia*, *Anemonastrum narcissiflorum* and *Crepis sibirica*) is also reported from the Feleacu Hills (Pop et al., 1962; Schur, 1859), where one of our reference profiles for the forest zone was located. Although some occurrences of rare species may be related to the local calcareous spring fens (Gałka et al., 2018; Tanţău and Fărcaş, 2001), it is possible that mesic open habitats may have existed for a long time also in this area, where forest vegetation clearly played a greater role during the Holocene than in the adjacent forest-steppe.

Representation of trees in the soil charcoal assemblages of Transylvanian forest-steppe grasslands is even lower than in analogous communities in the White Carpathian Mts in the Czech Republic, where we also reconstructed Holocene continuity of open to semi-open forest-steppe landscape (Hájková et al., 2011; Novák et al., 2019). The most similar is the Miliovy site at the foot of the White Carpathians, where steppe grasslands dominated by *Stipa tirsa* occurred in the past (Podpěra, 1930; Willner et al., 2019), i.e. the same community that is found today at Valea Florilor and Fânaţele Clujului I-Valea lui Craiu sites. A striking difference between both regions is the abundance of *Quercus*. In Transylvania it is almost absent in soil charcoal assemblages and also in the present landscape it is mostly confined to small forests, usually situated on northern slopes. On the other hand, *Quercus* is a major component of the White Carpathian forest-steppe (Doležal et al., 2010; Slachová, 2023) and it is also present, and in places abundant, in local soil charcoal assemblages (Novák et al., 2019). This difference is all the more striking because oak is one of the dominant species in the communities reconstructed as potential natural vegetation in both regions (Bohn et al., 2000–2003). Although its abundance in the pollen records from the Transylvanian forest-steppe is considerable (Feurdean et al., 2007a, 2015; Tantau et al., 2006), and oak forests were certainly part of the local vegetation mosaics, these records come areas that are also today more forested than the core part of the forest-steppe region between Cluj, Târgu Mureș and Ocna Mureș (Bădărău, 2005; Oancea and Valcea, 1987). Moreover, both profiles probably also capture pollen from the surrounding forested mountains, either due to their proximity (Avrig profile; Tantau et al., 2006), or due to the large source area of pollen at a lacustrine site (Lake Stiucii; Feurdean et al., 2015). New palynological data from the core part of the forest-steppe region could help to resolve this issue, however preservation of suitable sediment is a problem due to periodic drying of most of the local wetlands.

Another important topic for future studies is to clarify the role of woody plants in soil evolution, which appears to proceed in a non-linear and sometimes polygenetic way. Püntener et al. (2023) pointed out that chernozem-like soils under Transylvanian forest-steppe grasslands bear traces of woody plant occurrence (e.g. root channels). It is therefore necessary to determine whether these traces come from temperate trees missing from the soil charcoal record, or from the shrubs recorded. The analysis of n-alkanes and the C and N isotopic signal in the soil organic matter could also help in this respect, as shown for the White Carpathian forest-steppe by Karimi Nezhad et al. (under review). Further, given the great age of these soils (between the Last Glacial Maximum and Eemian Interglacial; Püntener et al., 2023), it is also possible that the woody plant traces come from the trees that once grew in the forests of the Last Glacial period (see discussion on these forests below).

### A prominent role of Juniperus

In soil charcoal analysis, it is doubly true that absence of evidence is not evidence of absence (Robin et al., 2013): if trees grew on the sites of present-day forest-steppe grasslands but did not burn, their charred remains could not be preserved. Therefore, positive evidence of the long-term open character of the forest-steppe grassland habitats is also important. We assume that the most consistent positive evidence in our dataset is the regular occurrence of *Juniperus.* This light-demanding coniferous shrub (particularly its most widespread subspecies *J. communis* subsp. *communis*) is considered an indicator of grazing, or rather successional stages after the retreat of intensive grazing. It favours a combination of soil disturbance and release of grazing pressure (Broome et al., 2017; Fitter and Jennings, 1975), while tolerating less intense fires (Diotte and Bergeron, 1989; Fitter and Jennings, 1975). It is rare in present-day forest-steppe grasslands, probably due to their unsuitable management in the past centuries (mowing). Nevertheless, it is a precious taxon in terms of their palaeoecological reconstruction, as it can be identified in both the charcoal and pollen records and indicates well the presence of open ecosystems (Lischke et al., 2013).

In most of the sites studied, *Juniperus* was the dominant species of the soil charcoal assemblages in deeper soil layers (below 1 m), largely corresponding to Late Glacial according to radiocarbon dating. It was often accompanied by *Pinus*. This agrees with the findings of palynological studies, where *Juniperus* was identified as one of the first woody species to spread during interglacial and interstadial warmings, shortly before or simultaneously with the expansion of *Pinus*, *Betula* and other pioneer woody species (Lischke et al., 2013; Reille et al., 2000). This indicates that *Juniperus* was an integral component of the Late Glacial vegetation on mesic sites in Transylvania. More importantly for the hypothesis being tested, *Juniperus* was also recorded in the upper soil layers of forest-steppe meadows, corresponding to the Holocene according to radiocarbon dating. We documented its occurrence during the Early (10,940 cal kyr BP, Valea lui Craiu; 8350 cal kyr BP, Valea Florilor), Middle (5400 cal kyr BP, Valea Caldă; 4690 cal kyr BP, Valea lui Craiu; 4520 cal kyr BP, Valea Florilor) and Late Holocene (600 cal kyr BP, Valea Florilor). Particularly at Valea Florilor and Boj-Cătun sites its presence along the profiles appears uninterrupted. The important role of *Juniperus* in some parts of the Transylvanian forest-steppe is also supported by the results of palynological studies. While in profiles from other parts of Transylvania *Juniperus* is either absent or only sporadically represented during the Holocene (Diaconeasa and Mitroescu, 1987; Feurdean et al., 2007a; Tantau et al., 2006), it forms an almost continuous curve in the forest-steppe region between the cities of Cluj and Dej (Feurdean et al., 2015). It is also represented in the soil charcoal record of analogous ecosystems in the White Carpathians (Novák et al., 2019).

### Species-rich hemiboreal forest: a precursor to species-rich steppe grasslands?

One of the most interesting results is the abundant evidence of the woody species present on the sites of today’s grasslands during the Last Glacial period. In addition to the dominant *Juniperus*, *Pinus* is regularly present and, in some cases, abundant, and *Betula*, *Larix*/*Picea* and Pomoideae are represented in the deeper soil layers, radiocarbon-dated between 13,240 and 43,660 cal kyr BP. Charcoals of temperate trees are represented in deeper soil layers only in the forest grassland profile Lita. The inverse radiocarbon dates from this locality probably originate from translocated charcoals of younger origin. The higher bioturbation rate and consequent soil mixing under forest vegetation is well known from literature (e.g. Bobek et al., 2018; Šamonil et al., 2010).

Soil charcoal composition is consistent with the view that during the younger stages of the Last Glacial, Transylvanian Basin supported vegetation of boreal forest-steppe (prevailing during warmer and/or more humid phases) and periglacial steppe (prevailing during colder and/or drier phases; Feurdean et al., 2007b, 2015; Tantau et al., 2006). Although we are not able to reconstruct the full species composition on mesic sites, in line with the available biogeographical evidence we assume that they were close to present-day South Siberian hemiboreal forests and related herb-rich steppe grasslands (Bădărău, 2005; Chytrý et al., 2010; Kleopov, 1990; Nimis et al., 1994; Roleček, 2007). These communities share many floristic elements with forest-steppe grasslands in Transylvania and the whole peri-Carpathian region, as illustrated by the comparison in Table 3. Thus, their glacial precursors might have been the source of a number of species now rare or having disjunct occurrences in Transylvania (e.g. *Achillea impatiens*, *Adenophora liliifolia*, *Adonis volgensis*, *Crepis sibirica*, *Iris ruthenica*, *Pulsatilla patens*, *Serratula coronata*). Based on the above analogy, these species can be understood as relicts of previously widespread glacial communities (Bădărău, 2005; Pop et al., 1962; Roleček et al., 2014). Another important aspect of the South Siberian analogues is their high species richness (Chytrý et al., 2012), which may have contributed to the extreme fine-scale species richness of contemporary forest-steppe grasslands via the relict species pool (Roleček et al., 2014). In addition to the limited spread of shady temperate forests, Late Glacial and Holocene climate of the Transylvanian Basin (Figure S1) may have contributed to the persistence of glacial species pool. Younger Dryas, the final cold episode of the Last Glacial period (ca 12,900–11,700 cal kyr BP), was rather mild here and despite notable reduction in forest cover, both steppe and (hemi)boreal forest were preserved in lower altitudes (Feurdean et al., 2007a, 2015; Tantau et al., 2006). During the Holocene, the relatively high altitude of the basin (around 400 m a.s.l. on average; Sanders et al., 2002) might have limited the retreat of glacial elements and the spread of xerothermic sub-Mediterranean species. Thus, we have a seemingly paradoxical situation here: mesic sites, if not subjected to modern intensification of farming, may host more archaic communities than ecologically extreme sites on steep sunny slopes. Similar scenario was suggested for Eastern European steppes by Kleopov (1990).

**Table 3.** Frequencies of shared species of the peri-Carpathian forest-steppe grasslands on mesic sites (f_B-M_; *Brachypodio-Molinietum* association according to Willner et al., 2019) and South Siberian hemiboreal forests (f_B-B_; *Brachypodio-Betuletea* class according to Ermakov et al., 1991). Species with minimum frequency of 15 % in both communities are shown. Closely related vicariant species are separated by a slash.

| Species | $f_{B-M}$ | $f_{B-B}$ |
| --- | --- | --- |
| *Brachypodium pinnatum* s.lat. | 82 | 82 |
| *Dactylis glomerata* | 62 | 34 |
| *Achillea millefolium* aggr. | 79 | 33 |
| *Viola hirta* | 55 | 32 |
| *Vicia cracca* s.lat. | 30 | 51 |
| *Inula salicina* | 29 | 36 |
| *Poa pratensis* s.lat. | 44 | 29 |
| *Cruciata glabra/krylovii* | 26 | 23 |
| *Trifolium pratense* | 23 | 21 |
| *Galium boreale* | 19 | 84 |
| *Galium verum* | 67 | 19 |
| *Hypochaeris maculata* | 19 | 25 |
| *Filipendula vulgaris* | 63 | 18 |
| *Pulmonaria mollis* s.lat. | 18 | 73 |
| *Avenula pubescens* | 31 | 17 |
| *Origanum vulgare* | 16 | 25 |
| *Primula veris* | 38 | 16 |
| *Campanula glomerata* | 33 | 15 |
| *Sanguisorba officinalis* | 15 | 56 |

### Drivers of persistence of the Transylvanian forest-steppe

The origin of steppe grasslands and the past distribution of forests and open habitats in the Transylvanian Basin has long been a subject of controversy (Bădărău, 2005; Borza, 1936; Enculescu, 1924; Feurdean et al., 2015; Schur, 1859; Soó, 1927). One of the main sources of the different opinions was the discrepancy between the relatively mesic climate of the area and the rich occurrence of steppe species. Some authors based their reconstructions on the potential of the area for forest growth and supported their conclusions with evidence of forest species occurrence in areas that are currently forest-free (e.g. Borza, 1936; Soó, 1927). Others emphasized the abundance of non-forest species (often with disjunct distributional ranges), the significant representation of chernozem-like soils and other evidence of the long-term continuity of open habitats (e.g. Bădărău, 2005; Enculescu, 1924).

The results of recent studies suggest that both views can be reconciled if we take into consideration all major factors driving forests-steppe development, i.e. climate, topography, soil and disturbance regime (Chytrý, et al. 2022; Erdős et al., 2022). The synthesis of Feurdean et al. (2018), based in part on case studies from the Transylvanian Basin, proposed a general model of Holocene grassland persistence in the East-Central European lowlands. The model incorporates dry climate, fire and grazing by wild herbivores during the Early Holocene. For the subsequent wetter period of the Middle Holocene, it envisages a varied regime of disturbance (domestic herbivore grazing, crop cultivation, deforestation, deliberate burning) by Neolithic humans and subsequent cultures that followed without a critical bottleneck for light-demanding species. Natural factors such as shallow soils, disturbances by wild animals and natural fires also continued to play a role. While the authors of the model conceded the persistence of grasslands primarily on sites with low biomass productivity, our results support the continuity of grasslands on mesic sites as well. Given the link of Neolithic subsistence to productive soils (e.g. Kiosak and Matviishyna, 2023; Petrasch, 2020; Shennan, 2018), this concept seems plausible. In terms of biogeography and vegetation history, our findings do not contradict the idea of the local survival of species from the Late Glacial forest-steppe. However, spatial scaling of this continuity and a more precise understanding of the role of woody plants requires further focused study.

### Novelty of our findings

Recent empirical findings on the origin and long-term dynamics of extremely species-rich forest-steppe grasslands lend credence to authors, who developed similar ideas a century ago. While Soó (1927) emphasized the former wider distribution of forests in the Transylvanian Basin, he also accepted the idea of the survival of steppe species in extreme habitats such as landslides and saline soils. He also admitted the possibility of the survival of relicts of the previous steppe period in the places settled by humans, which is actually a variation on the classical *Steppenheidetheorie* (Gradmann, 1901, 1933) that has been appreciated in recent years (Pokorný et al., 2015).

## Conclusion

The long-term disturbance regime was identified as a crucial driving factor of forest-steppe ecosystems in Transylvania. It should become an essential component of any model explaining the past dynamics of East-Central European forest-steppe. Our results also support the hypothesis that forest as a climax community was not always present during postglacial vegetation development, even on favourable mesic sites in the lowlands of temperate Europe.

## Acknowledgements

We thank local experts Anna Szabó and Eszter Ruprecht for their help in selecting the sampling sites. Co-operation of Michal Hájek and our field trip crew (Stanislav Němejc, Pavel Daněk, Kristýna Hošková, Jakub Jaroš, Dario Püntener) is acknowledged. We also thank Anna Větvičková for her help in preparing the figures.

## Supplementary material

Supplemental material for this article is available online.

