## Supplemental Figure S1 for "Relationship between extreme species richness and Holocene persistence of forest-steppe grasslands in Transylvania, Romania"

Temperature [°C]

Precipitation [mm]

| Age [years] | Mean summer temperature | Mean winter temperature |
| --- | --- | --- |
| 50 | 18.4 | -2.8 |
| 150 | 18.2 | -2.8 |
| 250 | 17.9 | -3 |
| 350 | 17.9 | -2.9 |
| 450 | 18.3 | -2.8 |
| 550 | 18.4 | -3.3 |
| 650 | 18.1 | -3.5 |
| 750 | 18 | -3.8 |
| 850 | 18 | -3.5 |
| 950 | 18.4 | -3.3 |
| 1050 | 18.2 | -3.3 |
| 1150 | 18 | -3.2 |
| 1250 | 18.1 | -3.2 |
| 1350 | 17.9 | -3.3 |
| 1450 | 18.3 | -4.1 |
| 1550 | 18.3 | -2.9 |
| 1650 | 18.4 | -3 |
| 1750 | 18.4 | -2.7 |
| 1850 | 18.6 | -3.8 |
| 1950 | 18.6 | -4 |
| 2050 | 18.2 | -3.6 |
| 2150 | 18.3 | -3.6 |
| 2250 | 18.4 | -3.3 |
| 2350 | 18.4 | -3.3 |
| 2450 | 18.6 | -3 |
| 2550 | 18.8 | -4.2 |
| 2650 | 18.6 | -3.4 |
| 2750 | 18.8 | -3.8 |
| 2850 | 19.1 | -3.5 |
| 2950 | 19.1 | -3.5 |
| 3050 | 18.8 | -3 |
| 3150 | 19.1 | -3.9 |
| 3250 | 18.8 | -3.4 |
| 3350 | 18.7 | -3.6 |
| 3450 | 19 | -3.5 |
| 3550 | 19.3 | -3.8 |
| 3650 | 19 | -3.8 |
| 3750 | 19.1 | -4 |
| 3850 | 19.1 | -4 |
| 3950 | 18.8 | -3.5 |
| 4050 | 18.7 | -3.5 |
| 4150 | 18.9 | -3.9 |
| 4250 | 19.3 | -3.3 |
| 4350 | 19.4 | -3.7 |
| 4450 | 19.3 | -3.7 |
| 4550 | 19.3 | -3.1 |
| 4650 | 19.7 | -3.6 |
| 4750 | 19.5 | -3.9 |
| 4850 | 19.2 | -3.7 |
| 4950 | 19.3 | -3.7 |
| 5050 | 19.1 | -3.8 |
| 5150 | 19.3 | -4.2 |
| 5250 | 19.7 | -4 |
| 5350 | 19.6 | -4.1 |
| 5450 | 19.3 | -3.9 |
| 5550 | 19.6 | -3.8 |
| 5650 | 20 | -3.9 |
| 5750 | 19.6 | -3.7 |
| 5850 | 19.8 | -4.1 |
| 5950 | 19.6 | -4.2 |
| 6050 | 19.6 | -4.2 |
| 6150 | 19.9 | -4 |
| 6250 | 19.8 | -4.4 |
| 6350 | 19.7 | -4 |
| 6450 | 20.1 | -4.2 |
| 6550 | 19.8 | -4.1 |
| 6650 | 19.9 | -3.5 |
| 6750 | 20 | -3.7 |
| 6850 | 19.7 | -4.3 |
| 6950 | 19.9 | -4 |
| 7050 | 20.2 | -4.5 |
| 7150 | 20 | -4.7 |
| 7250 | 20 | -4.8 |
| 7350 | 20.1 | -4.4 |
| 7450 | 19.8 | -4.5 |
| 7550 | 20 | -4.4 |
| 7650 | 20.1 | -4.9 |
| 7750 | 20.7 | -5.5 |
| 7850 | 20.3 | -4.7 |
| 7950 | 20.3 | -5.1 |
| 8050 | 19.7 | -5.5 |
| 8150 | 19.6 | -5.4 |
| 8250 | 19.7 | -5.2 |
| 8350 | 20 | -5.8 |
| 8450 | 19.8 | -5.5 |
| 8550 | 20.1 | -5.6 |
| 8650 | 20.1 | -5.5 |
| 8750 | 20 | -5.3 |
| 8850 | 20 | -5.8 |
| 8950 | 20 | -5.2 |
| 9050 | 20.9 | -5.7 |
| 9150 | 20.4 | -4.9 |
| 9250 | 20.2 | -5.5 |
| 9350 | 20.3 | -5.6 |
| 9450 | 20.2 | -5.8 |
| 9550 | 20.3 | -5.6 |
| 9650 | 20.2 | -5.8 |
| 9750 | 19.6 | -6.3 |
| 9850 | 19.9 | -5.8 |
| 9950 | 19.8 | -6 |
| 10050 | 20.6 | -5.9 |
| 10150 | 20.8 | -5.9 |
| 10250 | 19.9 | -6.5 |
| 10350 | 20 | -6.5 |
| 10450 | 20.2 | -6.6 |
| 10550 | 20.3 | -6.3 |
| 10650 | 20.2 | -6.4 |
| 10750 | 19.9 | -6.3 |
| 10850 | 20.3 | -6.4 |
| 10950 | 20.1 | -7 |
| 11050 | 20 | -6.3 |
| 11150 | 19.9 | -7.1 |
| 11250 | 19.6 | -6.7 |
| 11350 | 19.4 | -7.2 |
| 11450 | 19.5 | -7.9 |
| 11550 | 19.1 | -8.2 |
| 11650 | 19.4 | -8.8 |
| 11750 | 19.1 | -8.9 |
| 11850 | 19.4 | -8.7 |
| 11950 | 19.4 | -9.1 |
| 12050 | 19.1 | -10.6 |
| 12150 | 19.4 | -11.5 |
| 12250 | 18.2 | -10.1 |
| 12350 | 18.7 | -10 |
| 12450 | 18.3 | -9.7 |
| 12550 | 18.4 | -8.8 |
| 12650 | 18.5 | -8.8 |
| 12750 | 18.8 | -7.9 |
| 12850 | 18.9 | -8.1 |
| 12950 | 19.3 | -8.4 |
| 13050 | 19 | -8 |
| 13150 | 19 | -8.1 |
| 13250 | 19.2 | -8.2 |
| 13350 | 19 | -8.7 |
| 13450 | 19.1 | -8.6 |
| 13550 | 19 | -8.6 |
| 13650 | 18.9 | -8.5 |
| 13750 | 18.6 | -8.5 |
| 13850 | 18 | -9 |
| 13950 | 17.8 | -8.2 |
| 14050 | 17.7 | -8.1 |

|  | Mean annual precipitation |
| --- | --- |
| 50 | 510 |
| 150 | 470 |
| 250 | 458 |
| 350 | 481 |
| 450 | 469 |
| 550 | 498 |
| 650 | 470 |
| 750 | 467 |
| 850 | 472 |
| 950 | 498 |
| 1050 | 523 |
| 1150 | 502 |
| 1250 | 478 |
| 1350 | 445 |
| 1450 | 444 |
| 1550 | 459 |
| 1650 | 433 |
| 1750 | 415 |
| 1850 | 420 |
| 1950 | 389 |
| 2050 | 403 |
| 2150 | 424 |
| 2250 | 413 |
| 2350 | 441 |
| 2450 | 425 |
| 2550 | 474 |
| 2650 | 456 |
| 2750 | 488 |
| 2850 | 488 |
| 2950 | 466 |
| 3050 | 471 |
| 3150 | 464 |
| 3250 | 455 |
| 3350 | 454 |
| 3450 | 438 |
| 3550 | 454 |
| 3650 | 432 |
| 3750 | 465 |
| 3850 | 452 |
| 3950 | 446 |
| 4050 | 440 |
| 4150 | 468 |
| 4250 | 483 |
| 4350 | 492 |
| 4450 | 468 |
| 4550 | 478 |
| 4650 | 500 |
| 4750 | 488 |
| 4850 | 495 |
| 4950 | 510 |
| 5050 | 458 |
| 5150 | 489 |
| 5250 | 487 |
| 5350 | 483 |
| 5450 | 469 |
| 5550 | 453 |
| 5650 | 464 |
| 5750 | 483 |
| 5850 | 502 |
| 5950 | 493 |
| 6050 | 486 |
| 6150 | 527 |
| 6250 | 551 |
| 6350 | 510 |
| 6450 | 578 |
| 6550 | 561 |
| 6650 | 585 |
| 6750 | 555 |
| 6850 | 543 |
| 6950 | 557 |
| 7050 | 573 |
| 7150 | 544 |
| 7250 | 579 |
| 7350 | 595 |
| 7450 | 638 |
| 7550 | 597 |
| 7650 | 606 |
| 7750 | 614 |
| 7850 | 580 |
| 7950 | 576 |
| 8050 | 606 |
| 8150 | 640 |
| 8250 | 636 |
| 8350 | 640 |
| 8450 | 603 |
| 8550 | 650 |
| 8650 | 620 |
| 8750 | 595 |
| 8850 | 565 |
| 8950 | 597 |
| 9050 | 606 |
| 9150 | 623 |
| 9250 | 643 |
| 9350 | 661 |
| 9450 | 628 |
| 9550 | 631 |
| 9650 | 613 |
| 9750 | 599 |
| 9850 | 603 |
| 9950 | 634 |
| 10050 | 602 |
| 10150 | 631 |
| 10250 | 597 |
| 10350 | 606 |
| 10450 | 638 |
| 10550 | 603 |
| 10650 | 647 |
| 10750 | 623 |
| 10850 | 587 |
| 10950 | 621 |
| 11050 | 688 |
| 11150 | 655 |
| 11250 | 649 |
| 11350 | 656 |
| 11450 | 637 |
| 11550 | 637 |
| 11650 | 646 |
| 11750 | 624 |
| 11850 | 611 |
| 11950 | 582 |
| 12050 | 562 |
| 12150 | 652 |
| 12250 | 653 |
| 12350 | 661 |
| 12450 | 618 |
| 12550 | 673 |
| 12650 | 608 |
| 12750 | 614 |
| 12850 | 628 |
| 12950 | 602 |
| 13050 | 601 |
| 13150 | 556 |
| 13250 | 586 |
| 13350 | 579 |
| 13450 | 563 |
| 13550 | 601 |
| 13650 | 588 |
| 13750 | 571 |
| 13850 | 563 |
| 13950 | 561 |
| 14050 | 585 |

**Figure S1.** Late Glacial and Holocene temperature and precipitation simulations for the study area (CCSM3; Collins et al., 2006; Divíšek, 2021).
